# *Xanthomonas pateli* sp. nov. and *X. hingoranii* sp. nov. reveal a community of non-pathogenic *Xanthomonas* species complex in citrus

**DOI:** 10.64898/2026.09.16.751628

**Authors:** Anushika Sharma, Prabhu B. Patil

**Affiliations:** Bacterial Genetics, Genomics and Evolution Laboratory, CSIR-Institute of Microbial Technology, Chandigarh, India; The Academy of Scientific and Innovative Research, Ghaziabad, India

**Author notes:** **Author for Correspondence:** Prabhu B. Patil, Senior Principal Scientist, Bacterial Genetics, Genomics and Evolution Laboratory, CSIR-Institute of Microbial Technology, Chandigarh, India.

**Keywords:** *Xanthomonas*, Citrus, Non-pathogenic, Phylogenomics, Taxonomy

## Abstract

The genus *Xanthomonas* comprises a phylogenetically and ecologically diverse group of bacteria, historically characterized by economically important plant-pathogenic members affecting a wide range of agricultural crops. However, an increasing number of non-pathogenic and endophytic lineages have been recognized within the genus, expanding understanding of its taxonomic and ecological breadth. While it has been known for the last 30 years that non-pathogenic Xanthomonas (NPX) are present in diverse plants, studies from our lab over the last decade on the rice microbiome have revealed the presence of multiple, diverse NPX species as a community associated with a particular plant host. In this study, we identify two novel citrus plant-associated NPX species and provide evidence for the existence of an NPX community composed of multiple, diverse species, as in rice. Biochemical analysis, along with phylogenomic and genome-based taxonomic investigations, revealed that these strains represent two novel species within the genus *Xanthomonas*, providing evidence for a citrus-associated NPX community composed of multiple, diverse species, analogous to that described in rice. The lack of a canonical type III secretion system, along with multiple antimicrobial biosynthetic loci, as in the rice NPX community, points to their importance in plant microbiomes. Hence, such systematic studies of the NPX community need to be extended to all other plants. Accordingly, we propose that strain LMG 8992^T^ (DSM =122509^T^) be classified as a novel species, *Xanthomonas pateli* sp. nov., and strain LMG 8993^T^ (DSM = 122444^T^) be classified as a novel species, *Xanthomonas hingoranii* sp. nov.

## Introduction

Over the past 150 years, the genus *Xanthomonas* has been primarily studied as a group of highly specialized plant-pathogenic bacteria. To date, more than 39 species have been described, comprising over 200 pathovars that infect plants according to the tissue and host specific manner (1). However, there are increasing reports of non-pathogenic *Xanthomonas* species isolated from healthy plants. Notably, the presence of a community consisting of multiple and diverse *Xanthomonas* species within a single host has been reported. For instance, our previous studies revealed the presence of at least four novel *Xanthomonas* species associated with rice plants, suggesting their potential role in maintaining a healthy rice microbiome (2-6). Furthermore, several of these species were shown to inhibit the growth of *Xanthomonas oryzae* pv. *oryzae*, which causes bacterial blight disease in rice (5-7). Importantly, one of these species was found to be vertically transmitted and to function as a keystone species, contributing to plant growth promotion and disease protection (8). Collectively, these non-pathogenic *Xanthomonas* (NPX) species represent an excellent model for studying host–pathogen– microbiome interactions. Therefore, it is important to systematically investigate *Xanthomonas* communities associated with other host plants.

*Xanthomonas citri* pv. *citri* is a major pathogen of citrus plants and causes citrus canker, one of the most economically significant plant diseases worldwide (9). Similar to observations in rice, in the present study, we report the existence of a diverse community of *Xanthomonas* species associated with healthy citrus plants. Two of these species have been previously described as *Xanthomonas imtechensis* and *Xanthomonas rydalmerensis* (6, 10). In addition, we identify two new species and propose the names *Xanthomonas pateli* and *Xanthomonas hingoranii*. Consistent with NPX species identified in rice, all citrus-associated species described here are non-pathogenic to their host plants (11). Genomic analyses revealed the absence of the Type 3 secretion system (T3SS) and its associated effector proteins, which are required for pathogenicity. Conversely, several antimicrobial biosynthesis gene loci are present underscores their potential role as key members of the citrus microbiome.

Overall, this study highlights the importance of exploring NPX communities across diverse plant hosts to better understand their adaptation, ecological roles, and evolutionary trajectories as commensal bacteria.

### Origin and isolation of citrus NPX strains

In 1989, four non-pathogenic *Xanthomonas* strains were isolated from asymptomatic *Citrus* plants in the USA: LMG 8993^T^ from *Citrus* cv. Ina 69, LMG 8992^T^ from *Citrus* cv. Z-8 WDS, LMG 9002 from *Citrus* cv. AA1, and LMG 8989 from *Citrus* cv. Ina YZ. Two of the strains, LMG 9002 and LMG 8989, have previously been reported as *Xanthomonas rydalmensis* and *Xanthomonas imtechensis*, respectively (6, 10, 11). For our study, these strains were obtained from the BCCM/LMG Culture Collection.

### Genome sequencing, assembly and annotation of NPX strains from citrus

To obtain high-quality genome sequence of citrus NPX strains, bacterial strains grown overnight in Nutrient Broth (NB) at 28 °C with shaking at 180 rpm were pelleted by centrifugation, and genomic DNA was extracted using the Quick-DNA™ Fungal/Bacterial Miniprep Kit (Zymo Research, USA) according to the manufacturer’s instructions. The quality and quantity of genomic DNA were determined using a NanoDrop 1000 spectrophotometer (Thermo Fisher Scientific, USA). The genomic DNA was sent to Strand Life Sciences (Bengaluru, India) for paired-end whole-genome sequencing using the Illumina NovaSeq platform.

The quality of the paired-end raw reads was assessed using FastQC v0.11.9 (https://qubeshub.org/resources/fastqc). Adapter sequences and low-quality reads were removed using Trim Galore v0.6.7 (12). The filtered reads were assembled de novo using SPAdes v3.14.1 (13) with default parameters. Assembly statistics were generated using QUAST v5.0.2 (14). Genome completeness and contamination were estimated using CheckM v1.2.3 (15). The assembled genomes were submitted to the National Center for Biotechnology Information (NCBI) GenBank database and annotated using the NCBI Prokaryotic Genome Annotation Pipeline (PGAP).

The range of NPX genome sizes was 4.8-5 Mb. LMG 8993^T^ has a genome of about 4.9 Mb, and strain LMG 8992^T^ has a genome size of about 5 Mb. It was estimated that the genomes of *Xanthomonas imtechensis* (LMG 8989) and *Xanthomonas rydalmerensis* (LMG 9002) were approximately 4.8 Mb and 4.9 Mb, respectively. The G+C content ranged from 66% to 69%, with LMG 8992^T^ showing a comparatively higher G+C content (69%) than LMG 8993^T^ (66%). Between 4,039 and 4151, putative coding sequences were found in each of the four genomes. All strains had high genome completeness (99.9–100%), low contamination (0–0.74%), and high coverage, indicating high-quality genome assemblies. Overall, the genomic features of LMG 8992^T^ and LMG 8993^T^ are consistent with members of the genus *Xanthomonas*. Detailed genome statistics, including metadata, accession numbers and assembly metrics, are provided in **(Table 1)**.

**Table 1:**
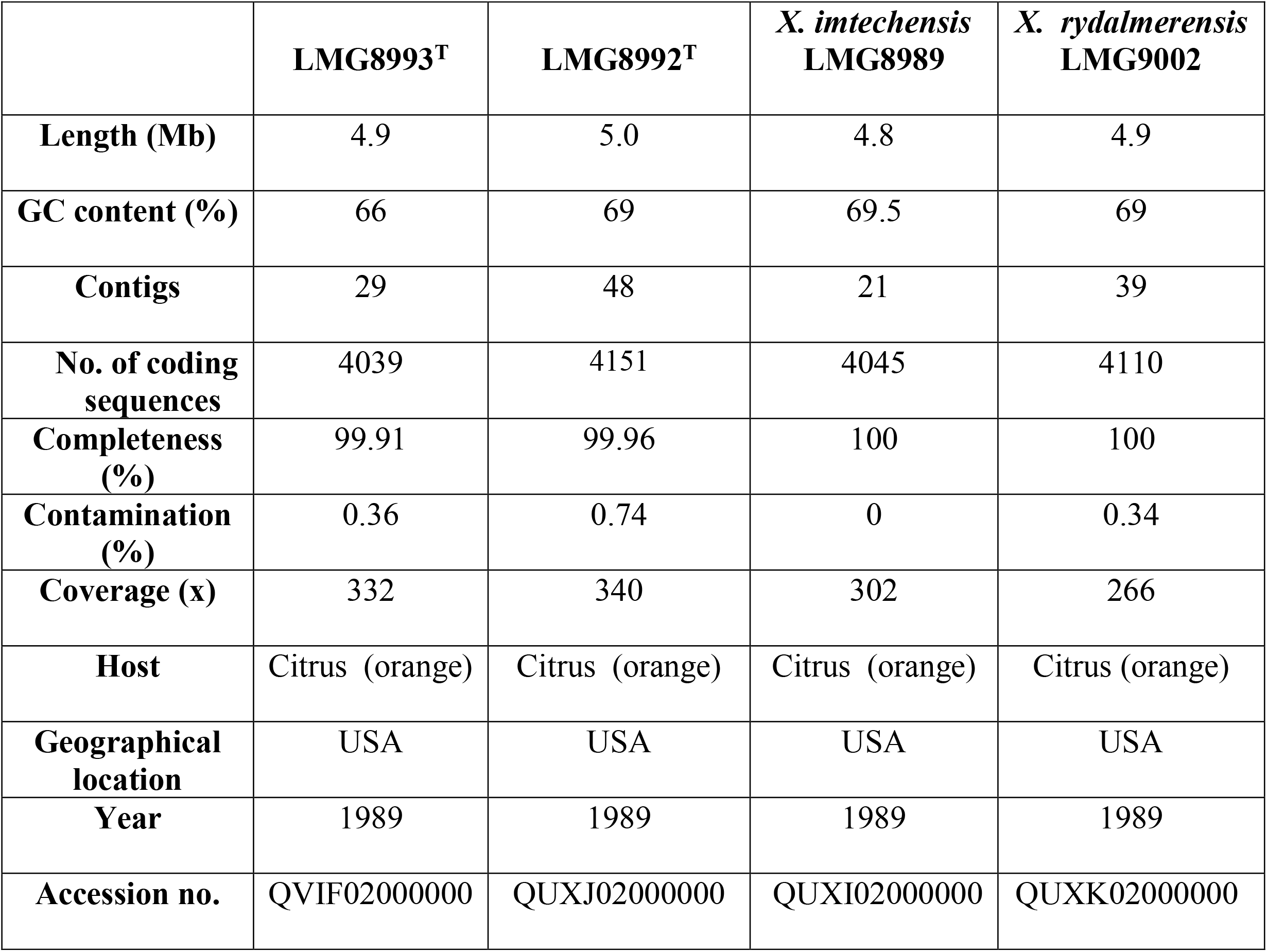
Genome assembly statistics and metadata of citrus-associated non-pathogenic *Xanthomonas* (NPX) community strains analyzed in this study are presented below.

### Genome-based phylogeny and taxonomy of citrus NPX strains

Complete 16S rRNA gene sequences were extracted from whole-genome assemblies of the novel strains and type strains of related *Xanthomonas* species using Barrnap v0.9 (https://github.com/tseemann/barrnap). For *X. maliensis* and *X. arboricola*, whose type-strain 16S sequences could not be extracted from available genome assemblies, sequences were instead obtained directly from NCBI. *Stenotrophomonas maltophilia* ATCC 13637^T^ was included as an outgroup to root the tree. Sequence alignment was performed in MUSCLE, followed by removal of poorly aligned or gap-containing regions using trimAl (default settings) (16, 17). A Neighbour-joining tree was subsequently reconstructed in MEGA 11.0.13 under the Jukes–Cantor substitution model, incorporating gamma-distributed among-site rate variation and evaluated with 1,000 bootstrap replicates (18). The resulting tree was visualized and annotated using iTOL v6 (19).

For the core gene-based phylogeny, genome assemblies of closely related *Xanthomonas* type strains were retrieved from NCBI. All genomes, including the novel strains and those obtained from NCBI, were re-annotated using Prokka (v1.14.6) (20), and core genes were identified using Roary (v3.13.0) (21), defined as those present in ≥99% of the genomes. Core-genome alignment was conducted with a minimum BLASTp identity of 90% and default parameters of the Markov Cluster Algorithm (MCL). RAxML v8.2.12 was used to construct a maximum-likelihood phylogenetic tree with the GTRGAMMA nucleotide substitution model and 1000 bootstrap replicates (22). iTOL was used to edit and view the final tree (19).

The majority of *Xanthomonas* species formed flat, poorly resolved clades in the 16S rRNA gene phylogeny, with many species not clearly separable from one another **(Supplementary Fig. 1)**. The 16S rRNA sequence of LMG 8992^T^ could not be differentiated from those of *X. rydalmerensis* and *X. sontii*, while that of LMG 8993^T^ showed no clear separation from *X. campestris* **(Supplementary Fig. 1)**. This lack of resolution is consistent with previous reports demonstrating that the 16S rRNA gene lacks sufficient discriminatory power for species-level classification within the genus *Xanthomonas* (23, 24). As core-genome phylogeny, ANI, and dDDH are required for accurate species delineation in this genus.

**Figure 1.**
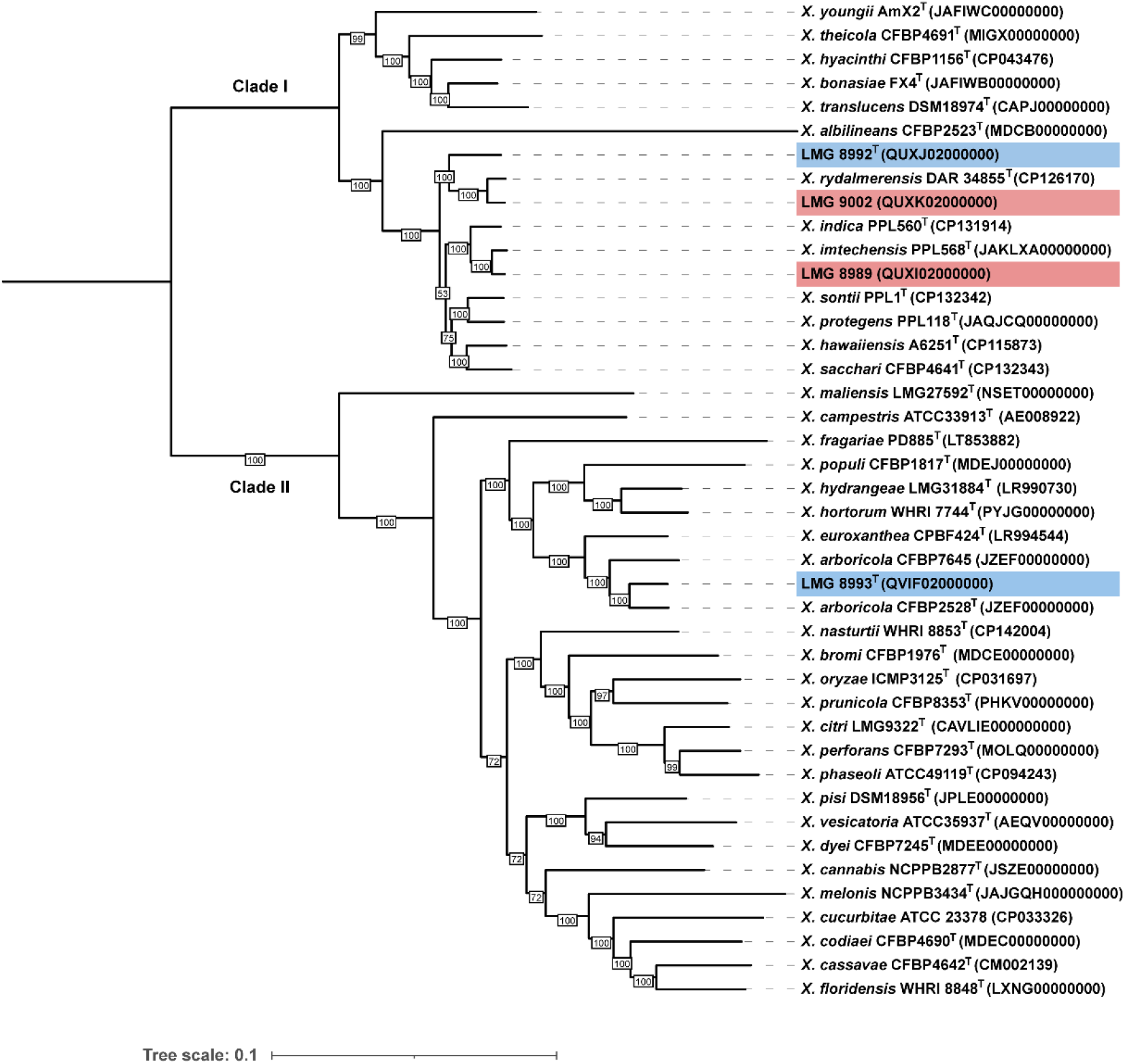
Using a representative of the genus *Xanthomonas*, a mid-rooted maximum likelihood tree was constructed using the core genes of the citrus-associated non-pathogenic *Xanthomonas*. Genome accession numbers are provided in parentheses after each strain name. The bootstrap values at the tree branches are greater than 50%. This study used strains that are indicated in blue. The number of nucleotide alterations per site is represented by the tree scale. The novel citrus-associated non-pathogenic *Xanthomonas* species described in this study are highlighted in blue, whereas the previously described citrus-associated non-pathogenic *Xanthomonas* species are highlighted in red.

Core-genome phylogeny placed the four citrus-associated *Xanthomonas* strains across both major clades of the genus. Three strains, LMG 8992^T^, LMG 9002, and LMG 8989, clustered within clade IB, consistent with clade IB representing the dominant lineage for non-pathogenic *Xanthomonas* (NPX) diversity, as previously reported for rice-associated NPX communities (7). LMG 8992^T^ formed a distinct cluster, most closely related to *X. rydalmerensis*. The fourth strain, LMG 8993^T^, fell within clade II, forming a separate cluster phylogenetically close to *X. arboricola* **(Fig. 1)**. Despite its phylogenetic proximity to *X. rydalmerensis*, LMG 8992^T^ was resolved as a distinct lineage, indicating novelty at the species level, as was LMG 8993^T^ within clade II. This distribution, with the majority of citrus-associated NPX species occurring in clade I, is reminiscent of NPX communities in rice, where most NPX species also fall within clade I, suggesting that clade I may serve as a dominant lineage for non-pathogenic *Xanthomonas* across plant hosts.

### Genome-based taxonomic indices further support the novel status of citrus NPX strains LMG 8992^T^ and LMG 8993^T^

To determine whether the citrus-associated isolates analyzed here match a previously identified *Xanthomonas* species, genome-relatedness indices were calculated. For this purpose, digital DNA–DNA hybridization (dDDH) using the Genome-to-Genome Distance Calculator 3.0 (formula 2), OrthoANI (v1.32) (25, 26) and ANIb calculated using JSpeciesWS were used to delineate species boundaries. The thresholds of ANI ≥95–96% and dDDH ≥70% were used to delineate species.

Based on the phylogenomic analyses, representative type strains from Clade I and Clade II of the genus *Xanthomonas* were selected for pairwise genomic comparisons using OrthoANI, ANIb and dDDH **(Table 2)**. To further evaluate the taxonomic placement of the isolates, the closest genomic relatives identified from these comparisons were subsequently analysed in detail, including their corresponding GGDC Model confidence intervals (CI), and are summarized in (**Supplementary Table S1)**. Pairwise comparisons were performed between the isolates LMG 8992^T^ and LMG 8993^T^, LMG 8989, and LMG 9002 against all type strains of validly described *Xanthomonas* species. All novel isolates showed ANI and dDDH values below the accepted species-level thresholds when compared to previously described strains. In addition, LMG 8993^T^ was most closely related to *X. arboricola* (OrthoANI/ANIb 96.2/95.8%, dDDH 67.2%, Model C.I. 64.2–70.0%), which is near the borderline for species delineation, while LMG 8992^T^ was closest to *X. rydalmerensis* (OrthoANI/ANIb 93.7/93.2%, dDDH 52.7– 51.6%, Model C.I. 50.1–55.4%). These values are below the species-level thresholds, confirming that LMG 8992^T^ and LMG 8993^T^ represent novel species **(Supplementary Table 1)**. LMG 9002 matched *X. rydalmerensis* (OrthoANI/ANIb 98.1/98.0%, dDDH 82.9%, Model C.I., 80.1–85.4%), and LMG 8989 corresponded to *X. imtechensis* (OrthoANI 98.3/98.1%, dDDH 84.9%, Model C.I., 82.2–87.3%). Taken together, the ANI and dDDH analyses, in combination with phylogenetic results, indicate that LMG 8992^T^ and LMG 8993^T^ represent novel species within the genus *Xanthomonas*, while LMG 8989 (*X. imtechensis*) and LMG 9002 (*X. rydalmerensis*) correspond to previously described species **(Table 2)**.

**Table 2:** Digital DNA–DNA hybridization (dDDH) and average nucleotide identity (OrthoANI/ANIb) values between the citrus NPX strains and representative type strains from the major phylogenetic clades of the genus *Xanthomonas*.

| S. No. | Strains | dDDH (in %) |  |  |  | OrthoANI/ANIb (in %) |  |  |  |
| --- | --- | --- | --- | --- | --- | --- | --- | --- | --- |
|  |  | LMG 8992<sup>T</sup> | LMG 8993<sup>T</sup> | X. *imtechensis* LMG 8989 | X. *rydalmerensis* LMG 9002 | LMG 8992<sup>T</sup> | LMG 8993<sup>T</sup> | X. *imtechensis* LMG 8989 | X. *rydalmerensis* LMG 9002 |
| 1. | LMG 8992<sup>T</sup> | 100 | 23.3 | 50 | 52.8 | 100.0/100 | 80.1/78.6 | 93.2/92.9 | 93.7/93.4 |
| 2. | LMG 8993<sup>T</sup> | 23.3 | 100 | 23.5 | 23.4 | 80.1/78.9 | 100.0/100 | 80.0/79.2 | 80.0/79.1 |
| 3. | LMG 8989 | 50 | 23.5 | 100 | 50.1 | 93.2/92.8 | 80.0/78.8 | 100.0/100 | 93.1/92.8 |
| 4. | LMG 9002 | 52.8 | 23.4 | 50.1 | 100 | 93.7/93.3 | 80.0/78.7 | 93.1/92.9 | 100.0/100 |
| 5. | *X. sontii* PPL1<sup>T</sup> | 51.6 | 23.4 | 50.1 | 50.6 | 93.5/92.9 | 79.9/78.7 | 93.1/92.8 | 93.2/92.9 |
| 6. | *X. sacchari* CFBP4641<sup>T</sup> | 49.8 | 23.3 | 51.2 | 48.8 | 93.0/92.9 | 79.9/78.7 | 93.6/93.1 | 92.8/92.5 |
| 7. | *X. indica* PPL560<sup>T</sup> | 49.8 | 23.4 | 68.5 | 49.8 | 93.0/92.8 | 79.9/78.9 | 96.3/96.2 | 93.1/92.8 |
| 8. | *X. imtechensis* PPL568<sup>T</sup> | 50.1 | 23.5 | 84.9 | 50.1 | 93.2/92.8 | 80.0/78.9 | 98.3/98.1 | 93.3/92.8 |
| 9. | *X. rydalmerensis* DAR34855<sup>T</sup> | 52.8 | 23.3 | 50.1 | 82.9 | 93.7/93.2 | 79.9/78.7 | 93.0/92.9 | 98.1/98.0 |
| 10. | *X. albilineans* CFBP2523<sup>T</sup> | 28.3 | 21.6 | 28.5 | 28.3 | 84.4/83.4 | 77.5/76.5 | 84.2/83.9 | 84.3/83.8 |
| 11. | *X. translucens*<br>DSM18974<sup>T</sup> | 32.9 | 24.3 | 32.9 | 32.9 | 87.2/86.0 | 80.3/79.0 | 87.0/86.3 | 87.1/86.2 |
| 12. | *X. bonasiae*<br>FX4<sup>T</sup> | 32.5 | 23.8 | 32.8 | 32.6 | 87.3/86.5 | 80.2/79.0 | 87.2/86.8 | 87.1/86.7 |
| 13. | *X. hyacinthi*<br>CFBP1156<sup>T</sup> | 33.3 | 23.7 | 33.3 | 33.4 | 87.6/86.7 | 80.3/79.1 | 87.5/86.8 | 87.4/86.8 |
| 14. | *X. hydrangeae*<br>LMG31884<sup>T</sup> | 23.0 | 38.9 | 23.0 | 23.0 | 79.3/78.3 | 89.7/89.3 | 79.4/78.3 | 79.2/78.4 |
| 15. | *X. theicola*<br>CFBP4691<sup>T</sup> | 32.1 | 23.6 | 32.4 | 32.3 | 86.7/85.3 | 80.2/78.8 | 86.8/86.1 | 86.7/85.6 |
| 16. | *X. youngii*<br>AmX2<sup>T</sup> | 30.7 | 23.9 | 30.6 | 30.5 | 86.1/85.1 | 80.4/79.2 | 85.9/85.3 | 86.2/85.2 |
| 17. | *X. maliensis*<br>LMG27592<sup>T</sup> | 22.9 | 26.7 | 23.0 | 22.9 | 79.4/78.3 | 83.1/82.4 | 79.5/78.8 | 79.5/78.6 |
| 18. | *X. arboricola*<br>CFBP2528<sup>T</sup> | 23.3 | 67.2 | 23.4 | 23.3 | 79.8/79.1 | 96.2/95.8 | 79.7/79.0 | 80.0/79.0 |
| 19. | *X. campestris*<br>ATCC33913<sup>T</sup> | 22.9 | 30.9 | 23 | 22.9 | 79.4/78.5 | 85.9/85.5 | 79.3/78.6 | 79.6/78.5 |
| 20. | *X. citri*<br>LMG9322<sup>T</sup> | 22.8 | 32.8 | 23.0 | 22.7 | 79.1/78.2 | 87.0/86.6 | 79.1/78.4 | 79.1/78.2 |
| 21. | *X. oryzae*<br>ICMP3125<sup>T</sup> | 22.9 | 32.5 | 22.9 | 22.7 | 78.9/78.0 | 86.8/86.1 | 78.7/78.1 | 78.9/78.1 |
| 22. | *X. euroxanthea*<br>CPBF424<sup>T</sup> | 23.4 | 48.3 | 23.4 | 23.4 | 80.2/79.1 | 92.7/92.3 | 79.8/79.2 | 80.2/79.2 |
| 23. | *X. vesicatoria*<br>ATCC35937<sup>T</sup> | 22.6 | 32.3 | 22.7 | 22.5 | 79.1/78.1 | 86.8/86.3 | 78.8/78.1 | 79.1/78.1 |

### Validation of novel species status of citrus NPX strains using TYGS

In addition, the genome sequences were uploaded to the Type (Strain) Genome Server (TYGS), which generated a whole-genome phylogeny using the d4 distance formula and the GBDP approach (27). In the TYGS analysis, LMG 8992^T^ and LMG 8993^T^ did not cluster with any previously described *Xanthomonas* species, supporting their novelty **(Fig. 2)**, while LMG 8989 and LMG 9002 corresponded to *Xanthomonas imtechensis* and *Xanthomonas rydalmerensis*, respectively (6, 10).

**Figure 2.**
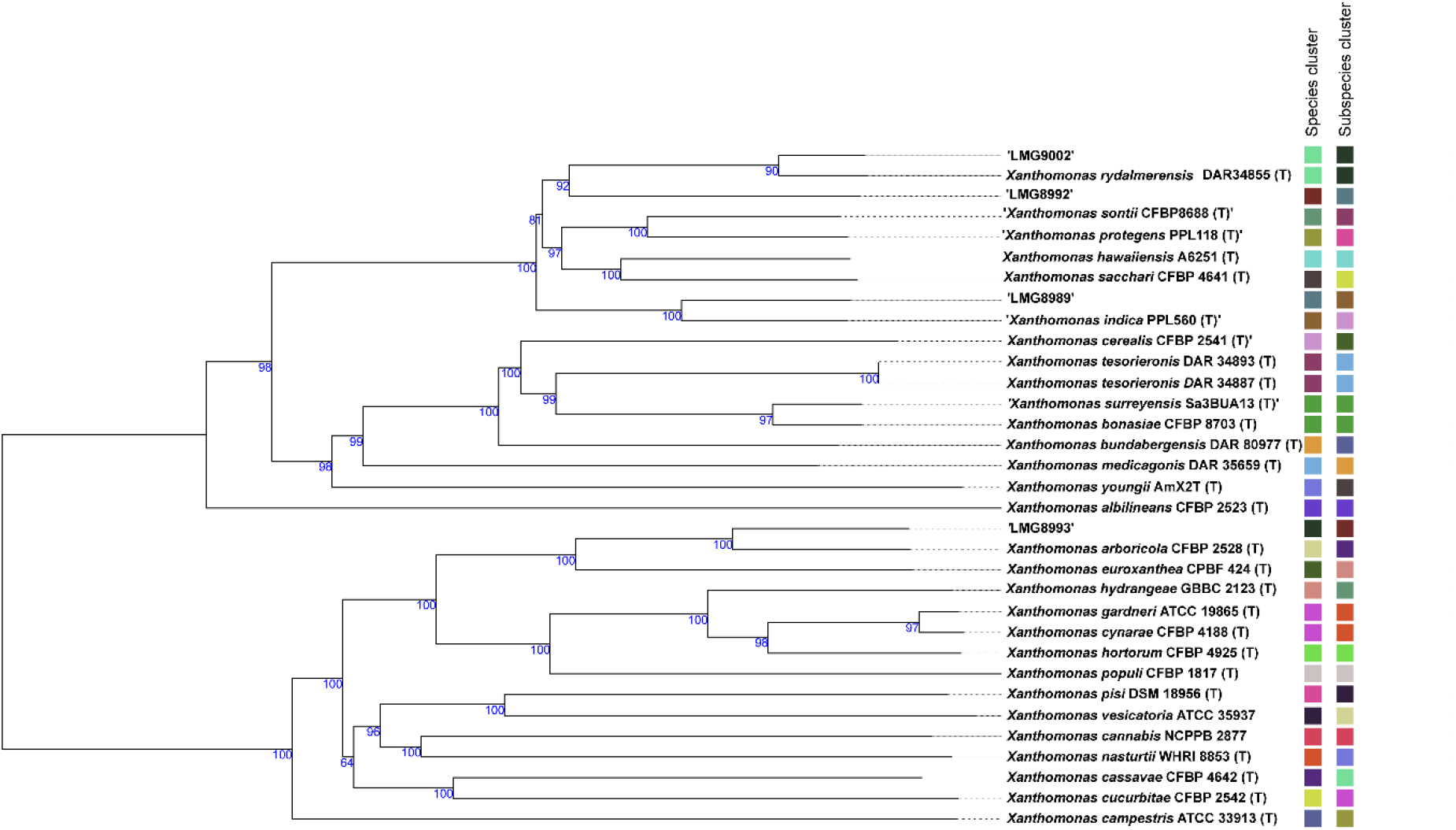
TYGS whole genome-based phylogeny of the non-pathogenic *Xanthomonas* from citrus using the database top hits. The colored bars in front of strains indicate different species clusters. (T) indicates the type strain of the corresponding species.

### Pangenome profile of citrus NPX strains

Core and accessory gene content across the novel *Xanthomonas* strains and closely related species was determined using a Roary-based pangenome analysis. Gene family counts were subsequently extracted and visualized as flower plots using custom scripts in RStudio (28), illustrating the distribution of core and unique genes. The flower diagrams **(Fig. 3a and 3b)** highlight the core gene shared among all strains at the centre, with petal sizes representing strain-specific unique genes. Each of the NPX species associated with citrus has a high number of unique genes supporting their distinct species status as established by genome-based phylogeny and taxonomy. LMG 8992^T^ and LMG 8993^T^ have high numbers of unique genes, 424 and 379, respectively, further supporting their novel species status. In fact, LMG 8992^T^ has the highest number of unique genes among related species in the clade.

**Figure 3.**
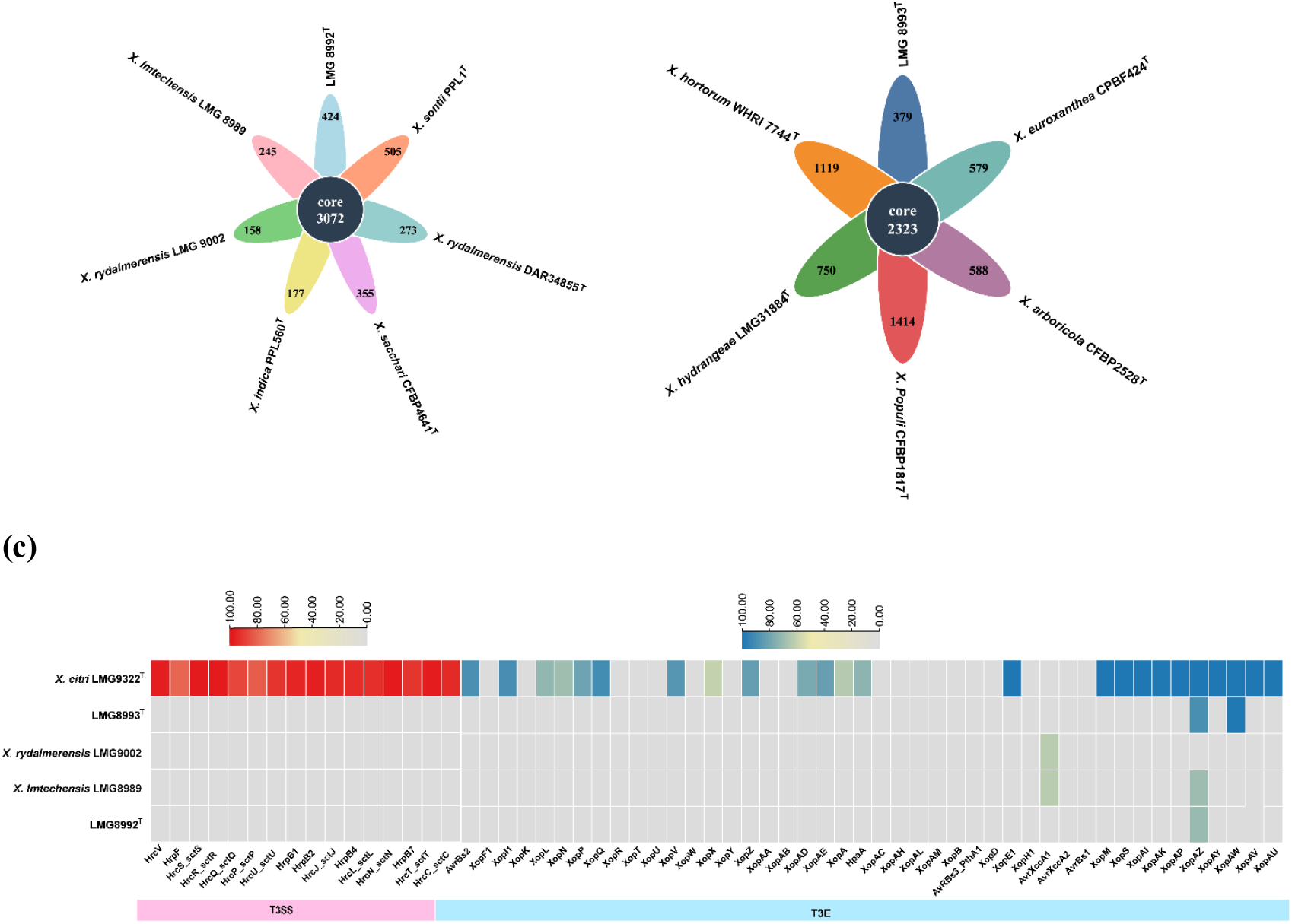
The floral plot displays **(a)** Clade 1 representative *Xanthomonas*, along with the novel strain LMG 8992^T^; the number of core genes in the centre and the number of unique genes for each strain are shown in the flower petals. **(b)** Clade 2 representative *Xanthomonas*, along with the novel strain LMG 8993^T^; the number of core genes in the centre and the number of unique genes for each strain are shown in the flower petals. **(c)** T3SS and T3E, showing a complete repertoire in pathogenic *Xanthomonas citri* versus absence in citrus-associated non-pathogenic *Xanthomonas* strains.

### Virulence Determinants and Pathogenicity Potential of citrus NPX species

The presence of virulence-related factors in citrus-associated *Xanthomonas* strains was investigated using genome-based approaches. To assess pathogenic potential, type III secretion system (T3SS) genes were retrieved from *Xanthomonas oryzae* pv. *oryzae* strain PXO99A, and type III effector (T3E) repertoires were obtained from the *Xanthomonas* resource database. These sequences were utilized as search terms in tBLASTn (v2.12.0+) against the genomes, using thresholds of ≥40% amino acid identity and ≥60% query coverage. Presence–absence patterns of T3SS genes and T3Es were visualized using TBtools (v1.108). The analysis revealed the citrus pathogen *Xanthomonas citri* encodes a functional T3SS (type III secretion system) and related effector genes (T3Es), which are essential for pathogenicity **(Fig. 3c)**. In contrast, these components were absent in the citrus-associated non-pathogenic *Xanthomonas* (NPX) strains, consistent with their non-pathogenic nature and previous reports (11).

### Bioprotection potential of citrus NPX species

AntiSMASH was used to predict secondary metabolite biosynthesis gene clusters (BGCs) (29), revealing a diverse repertoire of clusters across all four citrus-associated NPX strains. A conserved NI-siderophore cluster corresponding to xanthoferrin was identified in multiple strains, suggesting a shared role in iron acquisition. Comparative analysis showed that the strains differed in their BGC composition. Strain LMG 8989 (*Xanthomonas imtechensis*) harboured clusters associated with NI-siderophores, RiPP-like compounds, redox cofactors, arylpolyenes, and hydrogen cyanide. LMG 9002 (*Xanthomonas rydalmerensis*) exhibited a more diverse profile, including lassopeptides, NI-siderophores, NRPS, redox-cofactor clusters, hybrid NRPS–polyketide synthases, RRE-containing systems, arylpolyenes, and RiPP-like clusters. The novel strain LMG 8992^T^ displayed a rich repertoire of BGCs, including NRPS-like, RiPP-like, NI-siderophore, redox-cofactor, NRPS, hybrid NRPS–polyketide synthase, RRE-containing, arylpolyene, and lassopeptide clusters. In contrast, LMG 8993^T^ showed a comparatively simpler profile, comprising lassopeptide, NRPS, redox-cofactor, and NI-siderophore clusters **(Table 3)**. These biosynthetic gene clusters, repertoire diversity and distribution show that citrus-associated non-pathogenic *Xanthomonas* strains have the capacity to produce wide range of secondary metabolites that could improve plant health and act as biocontrol agents.

**Table 3:** Overview of Biosynthetic Gene Clusters Identified in Citrus-associated non-pathogenic *Xanthomonas* Strains.

| Strain | Major BGC Types Identified |
| --- | --- |
| **LMG 8989**<br>(*X. imtechensis*) | NI-siderophore, RiPP-like, redox-cofactor, arylpolyene, hydrogen cyanide |
| **LMG 9002**<br>(*X. rydalmerensis*) | Lasso peptide, NI-siderophore, NRPS, redox-cofactor, hybrid NRPS–PKS, RRE-containing, arylpolyene, RiPP-like |
| **LMG 8992<sup>T</sup>** | NRPS-like, RiPP-like, NI-siderophore, redox-cofactor, NRPS, hybrid NRPS–PKS, RRE-containing, arylpolyene, lasso peptide |
| **LMG 8993<sup>T</sup>** | Lasso peptide, NRPS, redox-cofactor, NI-siderophore |

### Morphological and Phenotypic characterization of citrus-associated novel non-pathogenic *Xanthomonas* species

Strains LMG 8992^T^ and LMG 8993^T^ were grown on nutrient agar and subsequently incubated in nutrient broth at 180 rpm, 28 °C, overnight. Cells were harvested by centrifugation (2000 rpm, 10 min), washed twice in phosphate-buffered saline (PBS), and resuspended in PBS. The suspension was applied to a carbon-coated grid for 15 min, negatively stained with 2% phosphotungstic acid for 30 s, air-dried, and examined by transmission electron microscopy at 200 kV. Using Biolog GEN III MicroPlates (Biolog), the biochemical characterisation of strains LMG 8992^T^ and LMG 8993^T^ was carried out according to the procedure described in (10).

Cells of strains LMG 8992^T^ and LMG 8993^T^ were rod-shaped and possessed a single polar flagellum, as observed by transmission electron microscopy **(Fig. 4a and 4b)**. Phenotypic traits of the citrus-associated *Xanthomonas* strains LMG 8992^T^ and LMG 8993^T^ utilized a broad range of carbon sources, including D-maltose, dextrin, D-trehalose, gentiobiose, D-cellobiose, D-melibiose, D-fructose, sucrose, D-glucose, D-mannose, D-galactose, L-fucose, and glycerol, hydrolysed gelatin, and metabolized the amino and organic acids L-alanine, L-aspartic acid, L-glutamic acid, L-serine, citric acid, D-malic acid, α-ketoglutaric acid, L-lactic acid, propionic acid, acetoacetic acid, and acetic acid. Both strains grew at pH 5–6 and tolerated 1% NaCl; LMG 8992^T^ additionally grew at 4% NaCl and showed variable growth at 8% NaCl, whereas LMG 8993^T^ did not grow above 1% NaCl. The two strains could be differentiated by several traits: LMG 8992^T^ utilized D-turanose, β-methyl-D-glucoside, D-salicin, and myo-inositol and was resistant to rifamycin SV, vancomycin, and sodium butyrate, whereas LMG 8993^T^ gave negative or variable results for these substrates and compounds. Full Biolog GEN III results for both strains are provided in **(Supplementary Table S2)**. A comparison of the biochemical characteristics between the two novel strains and their closely related species is presented in (Table 3); with reference phenotypic data for the related species obtained from previously published descriptions (2, 6, 10, 30-32). Since *Xanthomonas rydalmerensis* DAR34855^T^ and *Xanthomonas arboricola* pv. *juglandis* were the closest relatives of LMG 8992^T^ and LMG 8993^T^, respectively, based on phylogenomic analyses. LMG 8992^T^ differed from *X. rydalmerensis* DAR34855^T^ by its positive utilization of formic acid and variable reaction for D-mannitol. LMG 8993^T^ differed from *X. arboricola* pv. *juglandis* by positive reactions for sucrose, citric acid and formic acid, whereas the *X. arboricola* strain was negative for these substrates **(Table 4)**.

**Table 4:** Differential Biolog characteristics of strains LMG 8992^T^ and LMG 8993^T^ and closely related *Xanthomonas* species. Data for strains LMG 8992^T^ and LMG 8993^T^ were obtained in the present study. Data for *X. sontii, X. sacchari, X. euroxanthea, X. imtechensis* and *X. rydalmerensis* were compiled from their respective species descriptions (protologues). Data for *X. arboricola* were taken from the published Biolog characterization of *X. arboricola* pv. *juglandis*. Symbols: **+**, positive; **−**, negative; v, variable; BD, borderline reaction; NR, not reported.

| Test | LMG 8992<sup>T</sup> | LMG 8993<sup>T</sup> | *X. imtechensis*<br>PPL568<sup>T</sup> | *X. rydalmerensis*<br>DAR34855<sup>T</sup> | *X. sontii*<br>PPL1<sup>T</sup> | *X. sacchari*<br>CFBP4641<sup>T</sup> | *X. euroxanthea*<br>CPBF424<sup>T</sup> | *X. arboricola*<br>LMG747<sup>T</sup> |
| --- | --- | --- | --- | --- | --- | --- | --- | --- |
| Sucrose | + | + | + | + | + | + | + | – |
| Turanose | + | v | + | + | + | + | – | NR |
| Raffinose | – | – | – | – | – | – | – | NR |
| α-D-Lactose | + | v | + | + | + | + | NR | + |
| β-Methyl-D-glucoside | + | – | + | + | + | + | – | – |
| L-Fucose | + | + | + | + | + | + | + | + |
| L-Rhamnose | – | – | – | v | – | – | – | – |
| D-Mannitol | v | – | – | – | – | – | NR | – |
| D-Arabitol | – | – | – | – | – | – | – | – |
| Formic acid | + | + | + | – | v | + | + | + |
| Quinic acid | + | – | v | + | NR | + | – | – |
| Citric acid | + | + | + | + | + | + | + | – |
| Bromo-succinic acid | + | + | + | + | NR | + | + | + |
| Propionic acid | + | + | + | + | NR | + | + | + |
| Acetic acid | + | + | + | + | NR | + | + | + |
| L-Alanine | + | + | + | + | + | + | + | + |
| L-Aspartic acid | + | + | + | + | + | + | NR | + |
| L-Glutamic acid | + | + | + | + | + | + | + | + |

**Figure 4.**
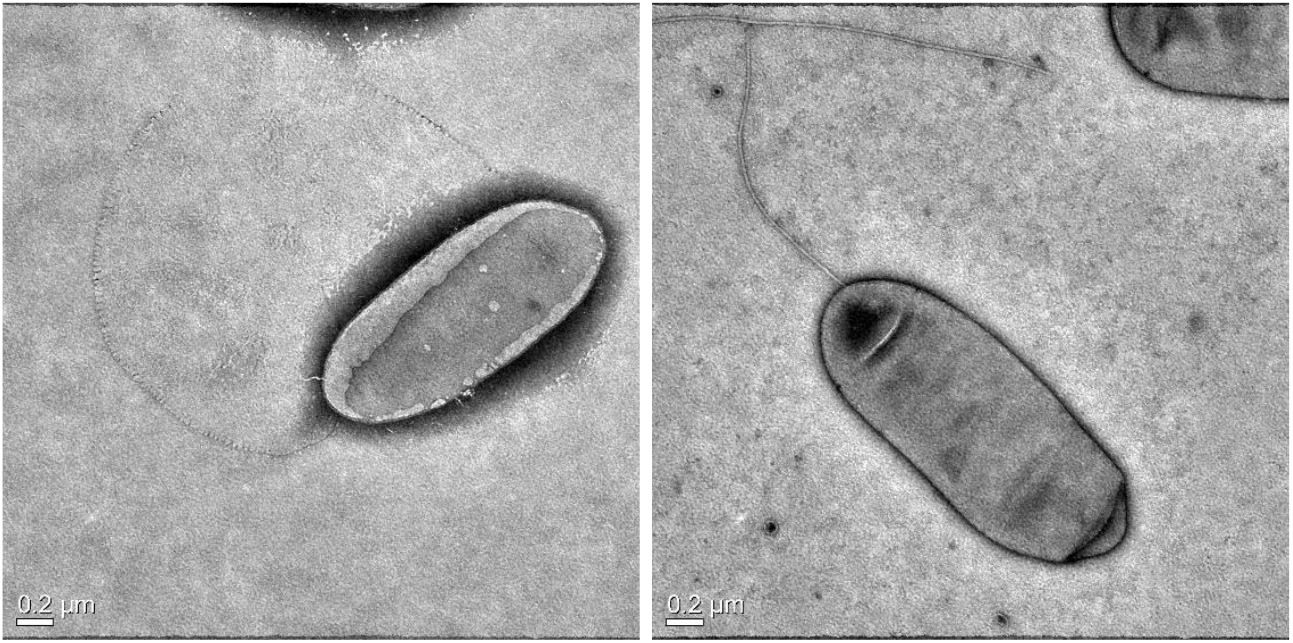
**(a)** Transmission electron micrographs of negatively stained cells of (a) Strain LMG 8992^T^ and **(b)** Strain LMG 8993^T^, showing rod-shaped cell morphology and a single polar flagellum. Scale bars, 0.2 µm.

### Description of *Xanthomonas pateli* sp. nov.

*Xanthomonas pateli* (pa.te’li. N.L. gen. n. *pateli*), of Patel, named in honour of Professor Makanji Kalyanji Patel, regarded as the father of phytobacteriology in India, in recognition of his foundational contributions to the field.

Cells of the type strain LMG 8992^T^ are rod-shaped, Gram-negative bacteria with a single polar flagellum. On nutrient agar, colonies are smooth, golden, and round after 48 hours of incubation at 28°C. This strain can withstand up to 4% NaCl, pH 5–6. Dextrin, D-maltose, D-trehalose, D-cellobiose, gentiobiose, sucrose, D-turanose, α-D-lactose, D-melibiose, and α-D-glucose are among the carbohydrates utilized. It also hydrolyses gelatin and pectin. L-glutamic acid, L-alanine, L-serine, citric acid, L-aspartic acid, D-malic acid, and L-histidine are among the organic acids and amino acids that it can use. Growth on 8% NaCl, and D-mannitol, sodium bromate was variable.

The type strain, LMG 8992^T^ = DSM 122509^T^, has a G+C content of 69% and an approximate genomic size of 5 Mb.

### Description of *Xanthomonas hingoranii* sp. nov.

*Xanthomonas hingoranii* (hin.go.ra’ni.i. N.L. gen. n. *hingoranii*), of Hingorani, named in honour of M. K. Hingorani (sic - M.K. Hingorani), pioneer of Indian agricultural microbiology, in recognition of his discovery of the bacterial etiology of pomegranate bacterial blight caused by a *Xanthomonas* species.

Cells of the type strain LMG 8993^T^ are Gram-negative, rod-shaped, non-spore-forming, and motile by means of a single polar flagellum. On nutrient agar, colonies are smooth, golden, and round after 48 hours of incubation at 28°C. Growth occurs at pH 5–6, and up to 1% NaCl. The strain utilizes dextrin, D-maltose, D-trehalose, D-cellobiose, gentiobiose, sucrose, D-melibiose, α-D-glucose, and N-acetyl-D-glucosamine, and hydrolyses gelatin and pectin. L-alanine, L-aspartic acid, L-glutamic acid, L-serine, citric acid, and D-malic acid are among the amino acids and organic acids metabolized. Growth on D-turanose, L-histidine, and D-gluconic acid was variable.

The type strain LMG 8993^T^ = DSM 122444^T^ has a G+C content of 66 % and a genomic size of about 4.9 Mb.

## Supporting information

Legends for Supplementary Figure and Tables

## Author contribution

A.S. designed the study, performed genome analyses, and drafted the manuscript. P.B.P. Conceived and participated in designing the study and finalizing the manuscript.

## Acknowledgements

We acknowledge the Department of Science and Technology - Innovation in Science Pursuit for Inspired Research (DST-INSPIRE) fellowship to A.S. The authors gratefully acknowledge Prof. Aharon Oren (The Hebrew University of Jerusalem, Jerusalem, Israel) for his valuable guidance on the nomenclature and etymology of the proposed species names.

## Author Notes

The whole-genome sequences obtained in this study have been deposited in the NCBI GenBank database under the following accession numbers: LMG 8992^T^, QUXJ02000000; LMG 8993^T^, QVIF02000000; *X. imtechensis* LMG 8989, QUXI02000000; and *X. rydalmerensis* LMG 9002, QUXK02000000. One supplementary figure and two supplementary tables are available with this article.

## Abbreviations

ANI: average nucleotide identity
ANIb: average nucleotide identity based on BLAST
BGC: biosynthetic gene cluster
dDDH: digital DNA–DNA hybridisation
NI-siderophore: non-ribosomal peptide synthetase-independent siderophore
NPX: non-pathogenic *Xanthomonas*
NRPS: non-ribosomal peptide synthetase
OrthoANI: orthologous average nucleotide identity
RiPP: ribosomally synthesized and post-translationally modified peptide
T3E: type III effector
T3SS: type III secretion system

## References

1. Parte AC, Sardà Carbasse J, Meier-Kolthoff JP, Reimer LC, Göker M. List of Prokaryotic names with Standing in Nomenclature (LPSN) moves to the DSMZ. International journal of systematic and evolutionary microbiology. 2020;70(11):5607–12.

2. Bansal K, Kaur A, Midha S, Kumar S, Korpole S, Patil PB. Xanthomonas sontii sp. nov., a non-pathogenic bacterium isolated from healthy basmati rice (Oryza sativa) seeds from India. Antonie Van Leeuwenhoek. 2021;114(11):1935–47.

3. Rana R, Madhavan VN, Saroha T, Bansal K, Kaur A, Sonti RV, et al. Xanthomonas indica sp. nov., a Novel Member of Non-Pathogenic Xanthomonas Community from Healthy Rice Seeds. Current Microbiology. 2022;79(10):304.

4. Triplett LR, Verdier V, Campillo T, Van Malderghem C, Cleenwerck I, Maes M, et al. Characterization of a novel clade of Xanthomonas isolated from rice leaves in Mali and proposal of Xanthomonas maliensis sp. nov. Antonie Van Leeuwenhoek. 2015;107(4):869–81.

5. Rana R, Sharma A, Madhavan VN, Korpole S, Sonti RV, Patel HK, et al. Xanthomonas protegens sp. nov., a novel rice seed-associated bacterium, provides in vivo protection against X. oryzae pv. oryzae, the bacterial leaf blight pathogen. FEMS Microbiology Letters. 2024;371:fnae093.

6. Sharma A, Patil PB. Xanthomonas imtechensis sp. nov.-a novel member of non-pathogenic Xanthomonas with bioprotection function from healthy rice seeds. bioRxiv. 2026:2026.02. 15.705894.

7. Rana R, Nayak PK, Madhavan VN, Sonti RV, Patel HK, Patil PB. Comparative genomics-based insights into Xanthomonas indica, a non-pathogenic species of healthy rice microbiome with bioprotection function. Appl Environ Microbiol. 2024;90(9):e0084824.

8. Rana R, Patil PB. Xanthomonas sontii, and Not X. sacchari, Is the Predominant Vertically Transmitted Core Rice Seed Endophyte. Phytopathology. 2024;114(9):2017–23.

9. Brunings AM, Gabriel DW. Xanthomonas citri: breaking the surface. Molecular plant pathology. 2003;4(3):141–57.

10. McKnight DJE, Wong-Bajracharya J, Okoh EB, Snijders F, Lidbetter F, Webster J, et al. Xanthomonas rydalmerensis sp. nov., a non-pathogenic member of Group 1 Xanthomonas. Int J Syst Evol Microbiol. 2024;74(3).

11. Vauterin L, Yang P, Alvarez A, Takikawa Y, Roth DA, Vidaver AK, et al. Identification of non-pathogenic Xanthomonas strains associated with plants. Systematic and applied microbiology. 1996;19(1):96–105.

12. Krueger F. Trim Galore!: A wrapper around Cutadapt and FastQC to consistently apply adapter and quality trimming to FastQ files, with extra functionality for RRBS data. Babraham Institute. 2015.

13. Prjibelski A, Antipov D, Meleshko D, Lapidus A, Korobeynikov A. Using SPAdes de novo assembler. Current protocols in bioinformatics. 2020;70(1):e102.

14. Gurevich A, Saveliev V, Vyahhi N, Tesler G. QUAST: quality assessment tool for genome assemblies. Bioinformatics. 2013;29(8):1072–5.

15. Parks DH, Imelfort M, Skennerton CT, Hugenholtz P, Tyson GW. CheckM: assessing the quality of microbial genomes recovered from isolates, single cells, and metagenomes. Genome Res. 2015;25(7):1043–55.

16. Capella-Gutiérrez S, Silla-Martínez JM, Gabaldón T. trimAl: a tool for automated alignment trimming in large-scale phylogenetic analyses. Bioinformatics. 2009;25(15):1972–3.

17. Edgar RC. MUSCLE: a multiple sequence alignment method with reduced time and space complexity. BMC bioinformatics. 2004;5(1):113.

18. Tamura K, Stecher G, Kumar S. MEGA11: Molecular Evolutionary Genetics Analysis Version 11. Molecular Biology and Evolution. 2021;38(7):3022–7.

19. Letunic I, Bork P. Interactive Tree of Life (iTOL) v6: recent updates to the phylogenetic tree display and annotation tool. Nucleic Acids Res. 2024;52(W1):W78–w82.

20. Seemann T. Prokka: rapid prokaryotic genome annotation. Bioinformatics. 2014;30(14):2068–9.

21. Page AJ, Cummins CA, Hunt M, Wong VK, Reuter S, Holden MT, et al. Roary: rapid large-scale prokaryote pan genome analysis. Bioinformatics. 2015;31(22):3691–3.

22. Stamatakis A. RAxML version 8: a tool for phylogenetic analysis and post-analysis of large phylogenies. Bioinformatics. 2014;30(9):1312–3.

23. Moore ERB, Krüger AS, Hauben L, Seal SE, De Baere R, De Wachter R, et al. 16S rRNA gene sequence analyses and inter- and intrageneric relationships of Xanthomonas species and Stenotrophomonas maltophilia. FEMS Microbiology Letters. 1997;151(2):145–53.

24. Hauben L, Vauterin L, Swings J, Moore E. Comparison of 16S ribosomal DNA sequences of all Xanthomonas species. International Journal of Systematic and Evolutionary Microbiology. 1997;47(2):328–35.

25. Lee I, Ouk Kim Y, Park S-C, Chun J. OrthoANI: an improved algorithm and software for calculating average nucleotide identity. International journal of systematic and evolutionary microbiology. 2016;66(2):1100–3.

26. Auch AF, von Jan M, Klenk H-P, Göker M. Digital DNA-DNA hybridization for microbial species delineation by means of genome-to-genome sequence comparison. Standards in genomic sciences. 2010;2:117–34.

27. Meier-Kolthoff JP, Göker M. TYGS is an automated high-throughput platform for state-of-the-art genome-based taxonomy. Nature communications. 2019;10(1):2182.

28. Team RC. R: A language and environment for statistical computing. R Foundation for Statistical Computing, Vienna, Austria. http://www.R-projectorg/. 2016.

29. Blin K, Shaw S, Vader L, Szenei J, Reitz ZL, Augustijn HE, et al. antiSMASH 8.0: extended gene cluster detection capabilities and analyses of chemistry, enzymology, and regulation. Nucleic Acids Res. 2025;53(W1):W32–w8.

30. Vauterin L, Hoste B, Kersters K, Swings J. Reclassification of xanthomonas. International Journal of Systematic and Evolutionary Microbiology. 1995;45(3):472–89.

31. Martins L, Fernandes C, Blom J, Dia NC, Pothier JF, Tavares F. Xanthomonas euroxanthea sp. nov., a new xanthomonad species including pathogenic and non-pathogenic strains of walnut. International Journal of Systematic and Evolutionary Microbiology. 2020;70(12):6024–31.

32. Kim HS, Cheon W, Lee Y, Kwon HT, Seo ST, Balaraju K, et al. Identification and Characterization of Xanthomonas arboricola pv. juglandis Causing Bacterial Blight of Walnuts in Korea. Plant Pathol J. 2021;37(2):137–51.

