## Supplementary material for "*Xanthomonas pateli* sp. nov. and *X. hingoranii* sp. nov. reveal a community of non-pathogenic *Xanthomonas* species complex in citrus": Legends for Supplementary Figure and Tables

**Supplementary Figure and Tables Legends:**

**Supplementary Fig. 1** Neighbour-joining phylogenetic tree based on 16S rRNA gene sequences, extracted from whole-genome assemblies using barrnap, showing the phylogenetic position of strains LMG 8992^T^ and LMG 8993^T^ among type strains of the genus Xanthomonas, with Stenotrophomonas maltophilia ATCC 13637^T^ as the outgroup. Branch colours indicate bootstrap support values as shown in the legend. Genome assembly accession numbers from which 16S rRNA sequences were extracted are given in parentheses. Bar, 0.01 substitutions per site.

**Supplementary Table S1:** Genomic relatedness of the citrus-associated non-pathogenic *Xanthomonas* strains LMG 8992^T^, LMG 8993^T^, LMG 8989 and LMG 9002 with their closest Xanthomonas type strains based on digital DNA–DNA hybridization (dDDH), Model confidence interval (CI), OrthoANI and ANIb analyses.

**Supplementary Table S2**: Full detailed Biolog GEN III characterization of LMG 8992^T^ and LMG 8993^T^.
