## Supplementary figures and images for "*Xanthomonas pateli* sp. nov. and *X. hingoranii* sp. nov. reveal a community of non-pathogenic *Xanthomonas* species complex in citrus"

### Supplementary Fig. 1

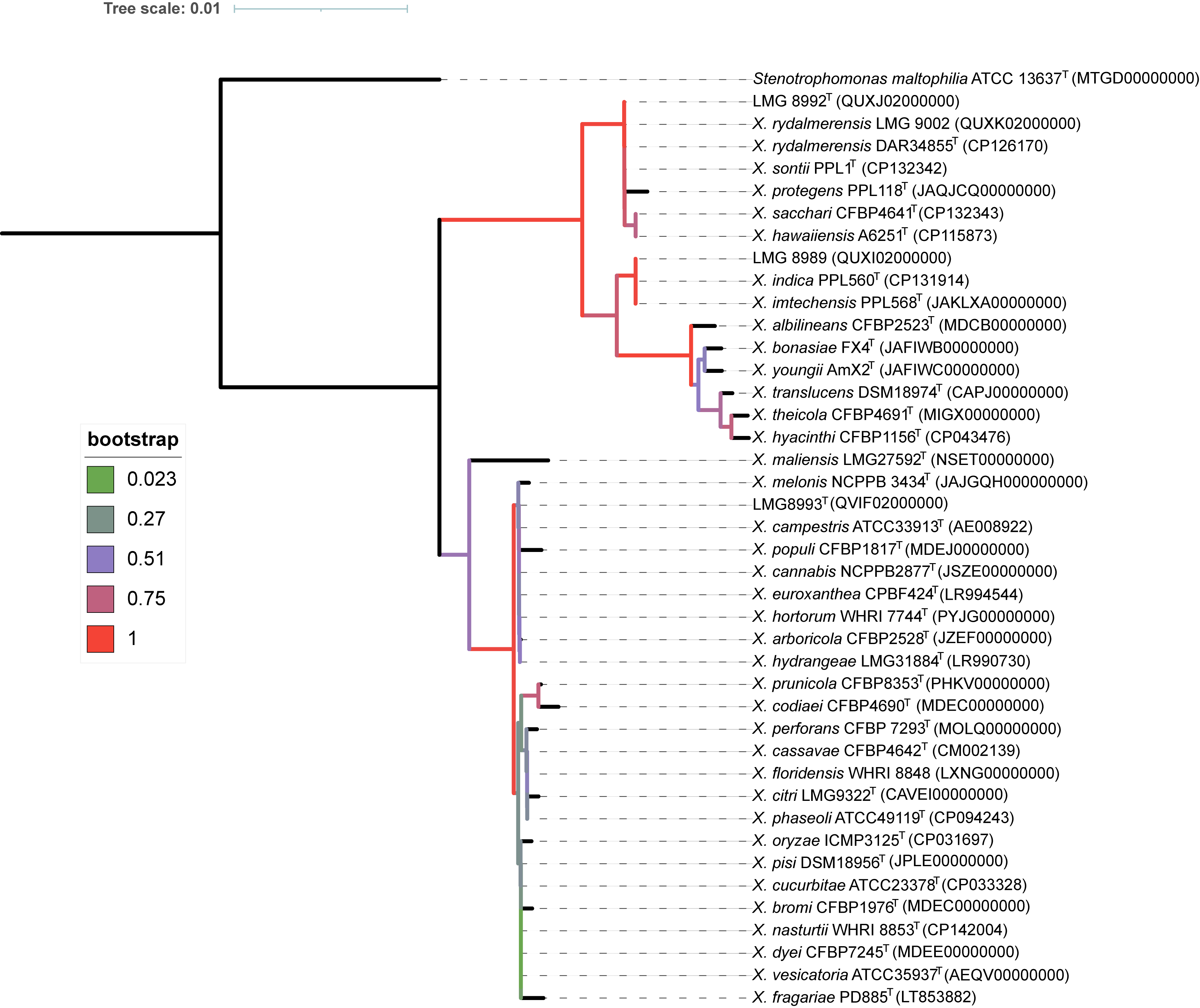
