## Supplementary Table S1 for "*Xanthomonas pateli* sp. nov. and *X. hingoranii* sp. nov. reveal a community of non-pathogenic *Xanthomonas* species complex in citrus"

| **Strain** | **Closest related strain** | **OrthoANI (%)** | **ANIb (%)** | **dDDH (%)** | **Model C.I. (%)** |
| --- | --- | --- | --- | --- | --- |
| **LMG 8992ᵀ** | X. rydalmerensis DAR34855ᵀ | 93.7 | 93.2 | 52.8 | **50.1–55.4** |
|  | LMG 9002 | 93.7 | 93.3 | 52.8 | **50.1–55.5** |
|  | LMG 8989 | 93.2 | 92.9 | 50.0 | **47.4–52.6** |
| **LMG 8993ᵀ** | X. arboricola CFBP2528ᵀ | 96.2 | 95.8 | 67.2 | **64.2–70.0** |
|  | X. euroxanthea CPBF424ᵀ | 92.7 | 92.3 | 48.3 | **45.7–50.9** |
| **LMG 8989** | X. imtechensis PPL568ᵀ | 98.3 | 98.1 | 84.9 | **82.2–87.3** |
|  | X. indica PPL560ᵀ | 96.3 | 96.2 | 68.5 | **65.5–71.3** |
|  | X. sacchari CFBP4641ᵀ | 93.6 | 93.1 | 51.2 | **48.6–53.9** |
| **LMG 9002** | X. rydalmerensis  DAR34855ᵀ | 98.1 | 98.0 | 82.9 | **80.1–85.4** |
|  | LMG 8992ᵀ | 93.7 | 93.3 | 52.8 | **50.1–55.5** |
|  | LMG 8989 | 93.1 | 92.9 | 50.1 | **47.5–52.7** |

**Supplementary Table: S1** Genomic relatedness of the citrus associated non-pathogenic *Xanthomonas* strains LMG 8992^T^, LMG 8993^T^, LMG 8989 and LMG 9002 with their closest Xanthomonas type strains based on digital DNA–DNA hybridization (dDDH), Model confidence interval (CI), OrthoANI and ANIb analyses.
